# Positive effects of low LDL-C and statins on bone mineral density: an integrated epidemiological observation analysis and Mendelian Randomization study

**DOI:** 10.1101/531137

**Authors:** Gloria Hoi-Yee Li, Ching-Lung Cheung, Philip Chun-Ming Au, Kathryn Choon-Beng Tan, Ian Chi-Kei Wong, Pak-Chung Sham

**Affiliations:** Department of Pharmacology and Pharmacy; Centre for Genomic Sciences; Department of Medicine, Li Ka Shing Faculty of Medicine, The University of Hong Kong, Hong Kong; Research Department of Practice and Policy, School of Pharmacy, University College London, London, UK; Department of Psychiatry, Li Ka Shing Faculty of Medicine, The University of Hong Kong, Hong Kong

**Keywords:** LDL-C, statins, bone mineral density, fracture, coronary artery disease, mendelian randomization

## Abstract

**Background:** Low-density lipoprotein cholesterol (LDL-C) is suggested to play a role in osteoporosis but its association with bone metabolism remains unclear. Effects of LDL-C-lowering drugs on bone are also controversial. We aim to determine whether LDL-C is linked causally to BMD and assess the effects of LDL-C-lowering drugs on BMD.

**Methods:** Association between blood lipid levels and BMD was examined by epidemiological observation analyses in US representative cohort NHANES III (N=3,638) and Hong Kong Osteoporosis Study (HKOS; N=1,128). Two-sample Mendelian Randomization (MR), employing genetic data from GWAS of blood lipids (N=188,577), total body BMD (TB-BMD) (N=66,628) and estimated BMD (eBMD) (N=142,487), was performed to infer causality between blood lipids and BMD. Genetic proxies for LDL-C-lowering drugs were used to examine the drugs’ effects on BMD.

**Results:** In NHANES III cohort, each SD decrease in LDL-C was associated with 0.045 SD increase in femoral neck BMD (95% CI: 0.009 to 0.081; P=0.015). A similar increase in BMD was observed in HKOS at femoral neck and lumbar spine. In MR analysis, decrease in genetically predicted LDL-C was associated with increase in TB-BMD [estimate per SD decrease, 0.038 (95% CI: 0.002 to 0.074); P=0.038] and eBMD [0.076 (0.042 to 0.111); P=1.20×10^−5^]. Reduction of TB-BMD was causally associated with increased LDL-C [0.035 (0.033 to 0.066); P=0.034]. Statins’ LDL-C-lowering proxies were associated with increased TB-BMD [0.18 (0.044 to 0.316); P=9.600×10^−3^] and eBMD [0.143 (0.062 to 0.223); P=5.165×10^−4^].

**Conclusions:** Negative causal association exists between LDL-C level and BMD. Statins’ LDL-C-lowering effect increases BMD, suggesting its protective effect on bone.

## Introduction

Osteoporosis and atherosclerosis are two major causes of morbidity and mortality. Emerging evidences have suggested a link between osteoporosis and coronary artery disease (CAD), and lipid metabolism was involved in the progression of both diseases^1^. Low-density lipoprotein cholesterol (LDL-C) is a well-known causal factor for atherosclerotic cardiovascular disease^2^. Other than this, the causal relationship among bone mineral density (BMD), CAD and LDL-C is still largely unknown.

The relationship between LDL-C and BMD has been investigated by different studies but the results remained inconclusive. Positive^3 4^, null^5 6^ and inverse^7–10^ associations between LDL-C and BMD were reported. Similar inconsistent findings are also observed for the association between BMD and CAD. A recent meta-analysis demonstrated that individuals with low BMD had an increased risk of developing cardiovascular disease and CAD^11^. Yet, a Mendelian Randomization (MR) study showed that one standard deviation (SD) increase in genetically predicted estimated BMD (eBMD) was associated with 5% higher risk of CAD^12^.

LDL-C has been a key target for CAD interventions and statins are one of the most commonly used LDL-C-lowering drug classes for treating CAD. Besides CAD, statins were also suggested to influence bone health though the evidence was inconsistent^13–15^. Another LDL-C-lowering drug, ezetimibe, showed an inverse association with BMD, although the association was not statistically significant^16^. While different levels of benefits and adverse effects were observed for different LDL-C-lowering drugs^17^, their effects on bone health have yet been investigated.

According to Mendel’s laws of inheritance, genetic variants are inherited randomly and exert a life-long effect on phenotypes. They could be used as instrumental variables to infer causality which is free of biases from confounding and reverse causation commonly found in observational studies. This approach is known as MR. Univariable MR has several key assumptions (Figure 1). A well-conducted MR study should provide reliable evidence that is comparable to those provided by randomized clinical trials (RCTs). The reliability could be further improved by cross-validation with evidence from other study designs^18^.

**Figure 1.**
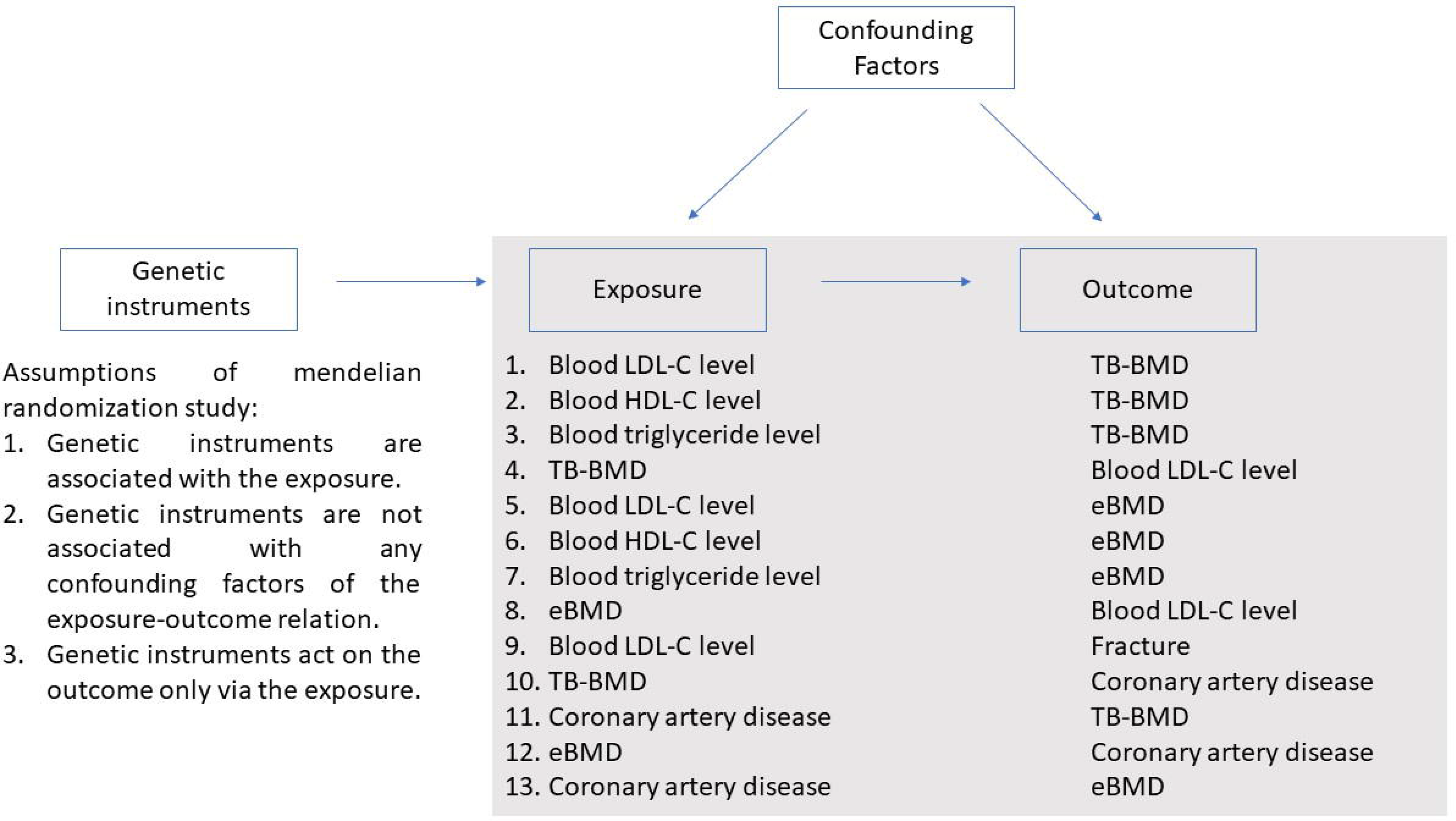
Key assumptions of univariable mendelian randomization and traits included in the MR analyses

Due to the inconsistent findings of previous observational studies, we firstly tested the association between LDL-C and BMD in two independent observational cohorts with distinct genetic compositions: the third National Health and Nutrition Examination Survey (NHANES III) and the Hong Kong Osteoporosis Study (HKOS) in the present study. Next, we determined genetic correlation and inferred causality between different traits using MR approach. We further tested if multiple LDL-C-lowering drugs (represented by genetic proxy) were associated with BMD variation.

## Materials and Methods

### Epidemiological observation analyses

Two independent cohorts of different genetic dispositions (NHANES III and HKOS) were employed in the epidemiological observational studies. For the analyses in the NHANES and HKOS, blood lipid levels and BMD were standardized with a mean of 0 and standard deviation (SD) of 1. The relationship between the blood lipid traits (including LDL-C, HDL-C and triglycerides) and BMD was evaluated using multivariable linear regression with adjustment for age, sex, ethnicity/race, height, weight, serum LDL-C levels (for the analysis with serum HDL-C and triglycerides levels), serum HDL-C levels (for the analysis with serum LDL-C and triglycerides levels), and serum triglyceride levels (for the analysis with serum HDL-C and LDL-C levels). Serum lipid levels were adjusted as covariates to avoid their potential pleiotropy with the lipid under investigation. For the NHANES analysis, sample weights that account for the unequal probabilities of selection, oversampling, and non-response were applied for all analyses using complex sampling module in SPSS version 22.0 software (SPSS Inc, Chicago, IL). All values presented were weighted to represent the U.S. civilian population. All statistical analyses were conducted using R or SPSS.

### Data sources for estimation of genetic correlation and MR

Summary statistics from large-scale genome-wide association study (GWAS) or meta-analysis of GWAS were used for both estimation of genetic correlation and MR. Two BMD phenotypes were assessed: total body BMD (TB-BMD) and eBMD at heel calcaneus as differences might exist between the two^9^. TB-BMD is measured by dual-energy X-ray absorptiometry (DXA), the gold-standard method of BMD measurement taking into account of both trabecular and cortical bone. Whereas, eBMD is estimated by quantitative ultrasound that primarily measures trabecular bone. Summary statistics for TB-BMD was obtained from the currently largest GWAS of DXA-derived BMD: a GWAS meta-analysis of 66,628 individuals from populations across America, Europe and Australia^19^ publicly available from GeFOS. Summary statistics for eBMD was obtained from a recent GWAS conducted in 142,487 participants of primarily European ancestry from UK Biobank, which was publicly available through the GEnetic Factors for OSteoporosis Consortium (GeFOS)^9^. While summary-level data obtained for lipids^20^, fracture^21^ and CAD^22^ were used in estimation of genetic correlation and evaluation of causal association, summary statistics of diabetes^23^ and BMI^24^ were utilized as confounding factors in MR analyses.

### Estimation of genetic correlation

LD score regression, which requires only GWAS summary statistics instead of individual-level data^25,26^, was employed to estimate the genetic correlation among LDL-C, TB-BMD, eBMD, fracture and CAD. Pre-computed LD scores suitable for European-ancestry samples and Python command line tool (http://github.com/bulik/ldsc) were adopted.

### Study design for MR and MR analyses

The current two-sample MR study utilized publicly available summary statistics from GWAS or GWAS meta-analysis: the instrument-risk factor and instrument-outcome associations were obtained from two different sets of participants (Figure 2). In brief, summary statistics of SNPs significantly associated with the risk factors were extracted from the confounding and outcome datasets. If the SNPs were not included in the datasets, proxies for the missing SNPs (r^2^>0.8) were identified from all the risk factors, confounding and outcome datasets. The SNPs were excluded from MR analyses if proxies were identified. Inverse-variance weighted (IVW)^27^ and multivariable IVW methods^28^ were used for main MR analysis. Weighted median^29^ and MR-Egger^30^ were used for sensitivity analysis. All MR analyses were conducted using the ‘MendelianRandomization’ package in R^31^. mRnd (http://cnsgenomics.com/shiny/mRnd/)^32^, an online web tool was employed to perform power calculation in our MR study. Power calculation and strength of genetic instruments are presented in Table 1.

**Table 1.**
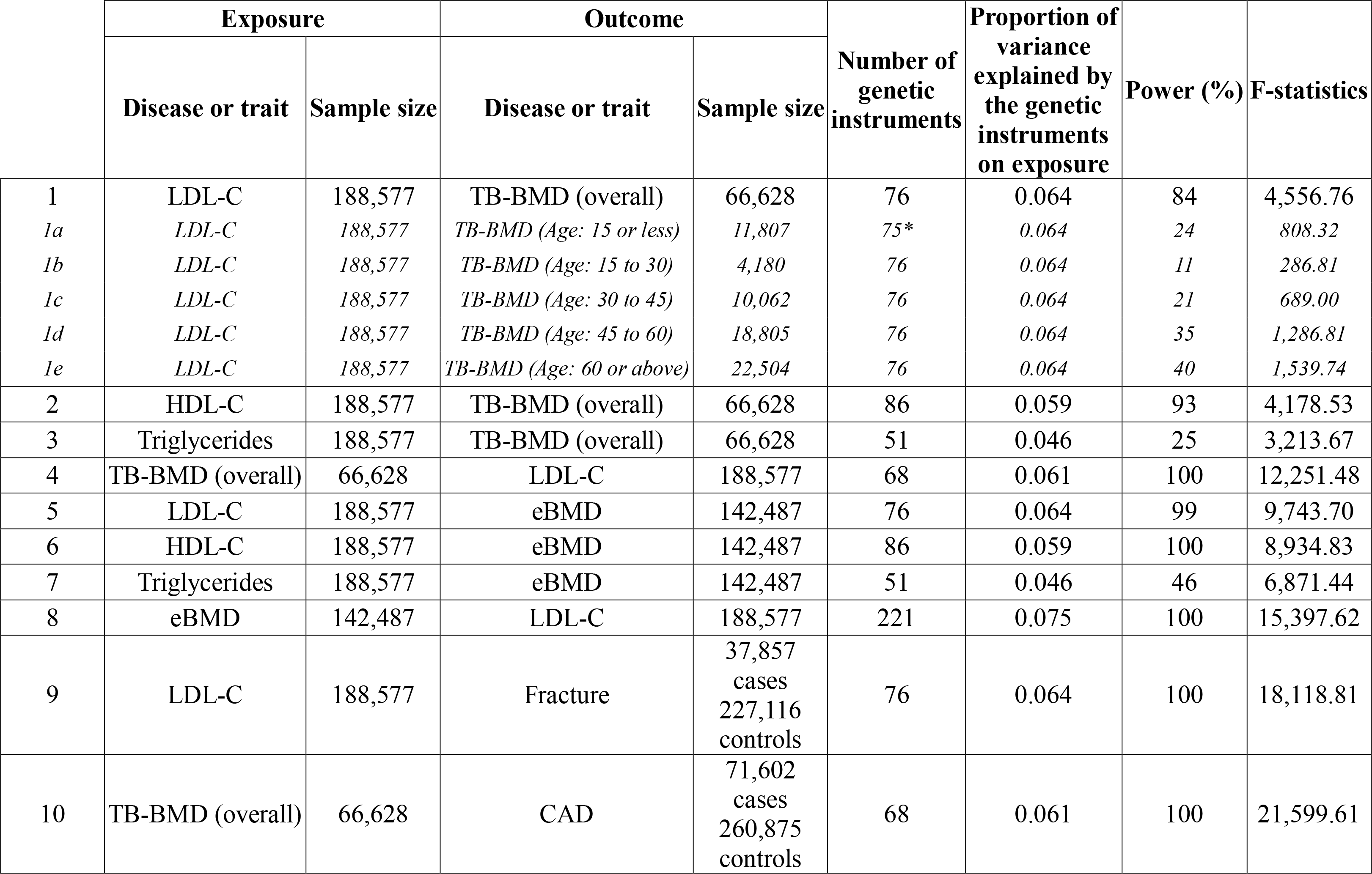
Power calculation for all the MR analyses conducted in the current study

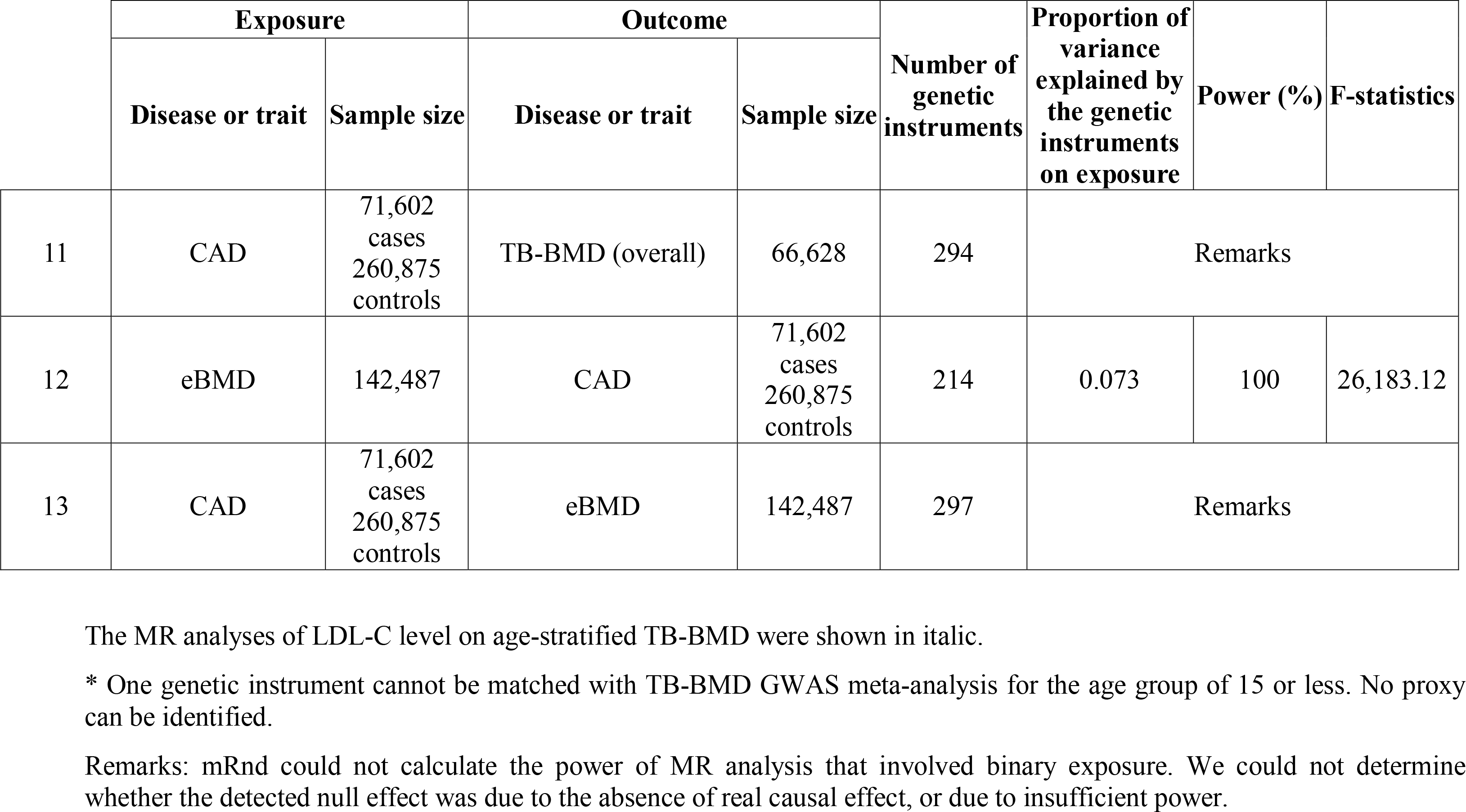

**Figure 2.**
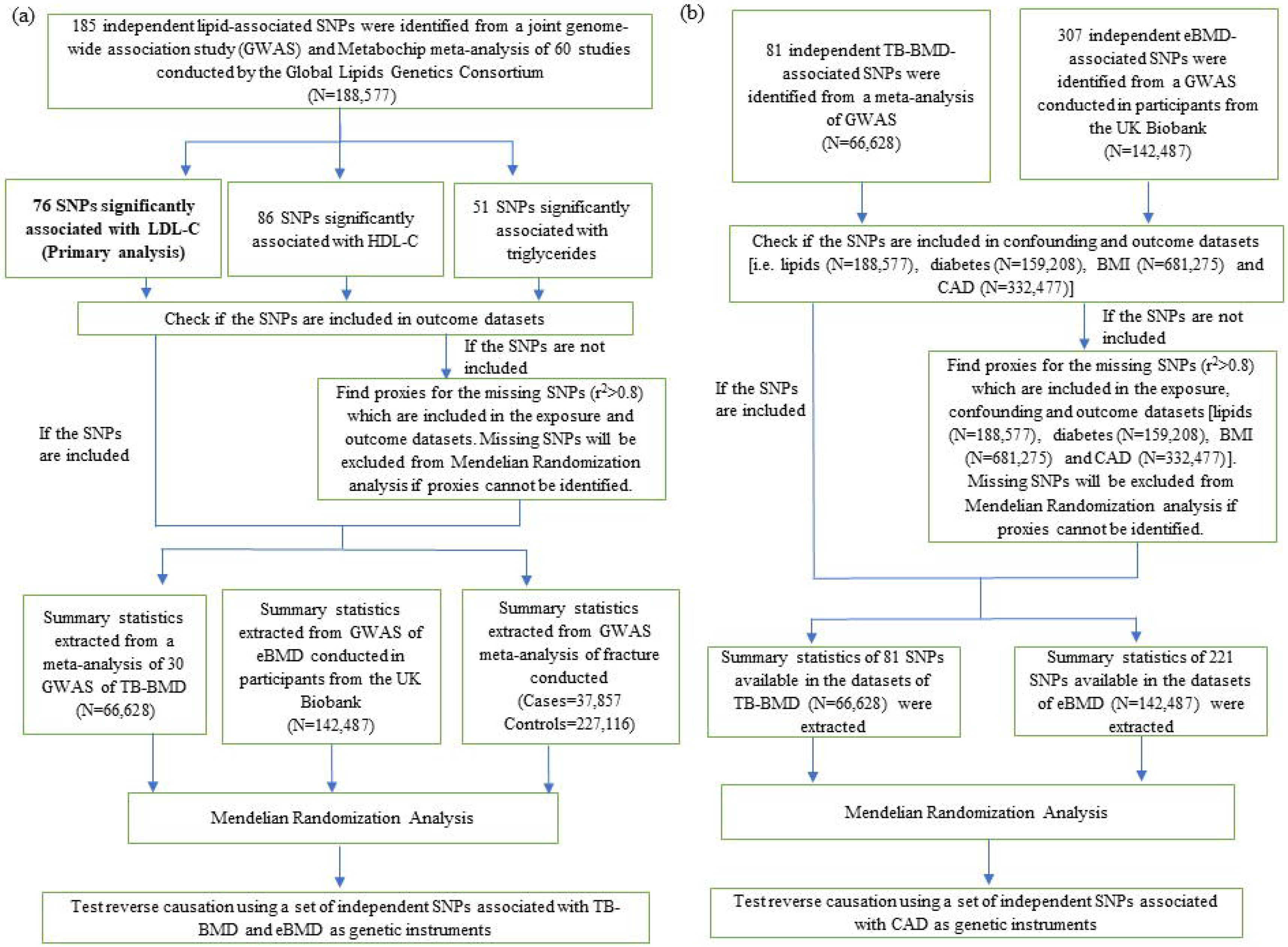
Study design and data sources adopted in meodeliao randomization study. (a) Evaluation of causality of blood lipid levels on BMD and frac.ture. (b) Evalua tion of causality of BMD on CAD.

### Drug target analysis using genetic proxy

Lotta *et al*. made use of LDL-C-lowering variants in or near genes encoding molecular targets of current or prospective LDL-C-lowering therapies as genetic proxies to study the efficacy of drugs on type 2 diabetes^33^. These genes included Niemann-Pick C1-Like 1 (*NPC1L1*) targeted by ezetimbe, 3-hydroxy-3-methylglutaryl-coenzyme A (*HMG-CoA*) reductase (*HMGCR*) targeted by statins, proprotein convertase subtilsin-kexin type 9 (*PCSK9*) targeted by Evolocumab and Alirocumab, the ATP-binding cassette subfamily G member 5 (*ABC5/ABC8*) targeted by bile acid sequestrants, and a prospective drug target low-density lipoprotein receptor (*LDLR*). In this study, we employed the same sets of SNPs as genetic instruments to examine if the LDL-C-lowering effects of the drugs would have causal association with TB-BMD and eBMD. Summary statistics were extracted from GLSC’s GWAS meta-analysis^34^, TB-BMD GWAS meta-analysis^19^ and eBMD GWAS^9^.

## Results

### Epidemiological observational analyses – Blood lipids and BMD

The association between serum lipids and BMD was evaluated in two epidemiological cohorts: the NHANES III and the HKOS. Demographic characteristics of participants from the two cohorts were shown in Table 2. The association between LDL-C and BMD was reported in Table 3. In the NHANES III cohort, each SD decrease of LDL-C was associated with 0.045 SD increase in BMD at femoral neck (95% CI: 0.009 to 0.081; P=0.015), after adjusting for age, sex, ethnicity/race, height, weight, serum HDL-C and triglyceride levels. A similar increase (0.039 SD) in BMD at femoral neck was also observed in the HKOS cohort but with a wider 95% CI (95% CI: −0.011 to 0.089; P=0.123). At lumbar spine, each SD decrease in LDL-C was associated with 0.083 SD increase in BMD (95% CI: 0.03 to 0.135; P=0.002). For HDL-C and triglycerides, no significant association was observed.

**Table 2.** Demographic characteristics of NHANES III and HKOS participants

|  | **NHANES III** |  |  | **HKOS** |  |  |
| --- | --- | --- | --- | --- | --- | --- |
| N | 3638 |  |  | 1128 |  |  |
| Age (years) | 57.1 | ± | 12.7 | 59.9 | ± | 10.5 |
| Female (%) | 1878 |  | 53.7% | 878 |  | 77.8% |
| Ethnicity/Race |  |  |  |  |  |  |
| Non-hispanic white (%) | 1917 |  | 80.8% | NA |  |  |
| Non-hispanic black (%) | 803 |  | 8.2% | NA |  |  |
| Mexican-American (%) | 766 |  | 3.2% | NA |  |  |
| Other (%) | 152 |  | 7.8% | NA |  |  |
| Southern Chinese (%) |  | NA |  | 1128 |  | 100% |
| Height (cm) | 167.4 | ± | 9.9 | 158.3 | ± | 8 |
| Weight (kg) | 76.3 | ± | 17.1 | 58.6 | ± | 10.6 |
| BMD at femoral neck (g/cm<sup>2</sup>) | 0.763 | ± | 0.147 | 0.7 | ± | 0.126 |
| BMD at lumbar spine (g/cm<sup>2</sup>) |  | NA |  | 0.925 | ± | 0.164 |
| BMI (kg/m<sup>2</sup>) | 27.15 | ± | 5.31 | 23.52 | ± | 6.34 |
| Serum LDL (mmol/L) | 3.53 | ± | 0.94 | 2.86 | ± | 0.81 |
| Serum HDL (mmol/L) | 1.31 | ± | 0.4 | 1.64 | ± | 0.46 |
| Serum triglycerides (mmol/L) | 1.6 | ± | 0.8 | 1.25 | ± | 0.66 |

Data are presented as mean±SD for continuous variables and N±%. for categorical variables.

**Table 3.** Comparison of observational and causal estimates for BMD (in SD) per 1 SD decrease of LDL-C

| Analysis | Beta | 95% CI |  | P-value |
| --- | --- | --- | --- | --- |
|  |  | Lower | Upper |  |
| **Observational** |  |  |  |  |
| BMD at femoral neck* |  |  |  |  |
| NHANES III | 0.045 | 0.009 | 0.081 | 0.015 |
| HKOS | 0.039 | -0.011 | 0.089 | 0.123 |
| BMD at lumbar spine* |  |  |  |  |
| HKOS | 0.083 | 0.03 | 0.135 | 0.002 |
| **Mendelian Randomization approach to evaluate causality of LDL-C on TB-BMD** |  |  |  |  |
| 76 SNPs |  |  |  |  |
| Conventional IVW | 0.038 | 0.002 | 0.074 | 0.038 |
| Multivariable IVW | 0.033 | -0.004 | 0.070 | 0.083 |
| Weighted median | 0.018 | -0.026 | 0.063 | 0.416 |
| MR-Egger | 0.011 | -0.045 | 0.068 | 0.694 |
| MR-Egger intercept | 0.002 | -0.001 | 0.005 | 0.235 |
| **Mendelian Randomization approach to evaluate causality of LDL-C on eBMD** |  |  |  |  |
| 76 SNPs |  |  |  |  |
| Conventional IVW | 0.076 | 0.042 | 0.111 | 1.20 x 10<sup>-5</sup> |
| Multivariable IVW | 0.083 | 0.047 | 0.118 | 4.39 x 10<sup>-6</sup> |
| Weighted median | 0.065 | 0.04 | 0.09 | 3.59 x 10<sup>-7</sup> |
| MR-Egger | 0.052 | -0.002 | 0.106 | 0.059 |
| MR-Egger intercept | 0.002 | -0.001 | 0.005 | 0.257 |

\*Model was adjusted for age, sex, ethnicity/race, weight, height, serum HDL-C levels, and serum triglycerides levels.

### Genetic correlation

Genetic correlation among the studied traits were shown in Table 4. Moderate and positive genetic correlation was observed between TB-BMD and eBMD (r=0.59; 95% CI: 0.555 to 0.628; P= 5.47 × 10^−59^). Weak and inverse genetic correlation was observed between LDL-C and both BMD phenotypes (TB-BMD: r=-0.079; 95% CI: −0.107 to −0.0519; P=0.0038; eBMD: r=-0.082; 95% CI: −0.1045 to −0.0597; P=0.0003). In addition, eBMD was weakly correlated with HDL-C (r=-0.0724; 95% CI: −0.0949 to −0.0499; P=0.0013) and CAD (r=0.0669; 95% CI: 0.0416 to 0.0922; P=0.0082). None of the remaining trait pairs had statistically significant genetic correlation.

**Table 4.** Genetic correlation among traits

|  | **Trait 1** | **Trait 2** | **Genetic correlation (95% CI)** | **P-value** |
| --- | --- | --- | --- | --- |
| 1 | TB-BMD | eBMD | 0.5918 (0.5553 to 0.6283) | 5.4657 x 10<sup>-59</sup> |
| 2 | LDL-C | TB-BMD | -0.0792 (-0.1065 to -0.0519) | 0.0038 |
| 3 | LDL-C | eBMD | -0.0821 (-0.1045 to -0.0597) | 0.0003 |
| 4 | HDL-C | TB-BMD | -0.0383 (-0.0616 to -0.015) | 0.0996 |
| 5 | HDL-C | eBMD | -0.0724 (-0.0949 to -0.0499) | 0.0013 |
| 6 | Triglycerides | TB-BMD | -0.0103 (-0.0377 to 0.0171) | 0.7085 |
| 7 | Triglycerides | eBMD | 0.0334 (0.0053 to 0.0615) | 0.2340 |
| 8 | TB-BMD | CAD | 0.0137 (-0.0166 to 0.044) | 0.6512 |
| 9 | eBMD | CAD | 0.0669 (0.0416 to 0.0922) | 0.0082 |
| 10 | LDL-C | Fracture | 0.024 (-0.0187 to 0.0667) | 0.5735 |

### MR analyses – blood lipids, TB-BMD and eBMD

The causal association between blood lipids and BMD were examined by MR. Primary analysis was done for LDL-C (Table 3). In IVW analysis, 1 SD decrease in genetically predicted blood LDL-C level was associated with 0.038 SD (95% CI: 0.002 to 0.074; P=0.038) increase in TB-BMD in IVW analysis. Similar estimate with a wider confidence interval was obtained for multivariable MR analysis adjusting for blood HDL-C and triglycerides levels, which are highly correlated with LDL-C. The result was no longer significant in sensitivity analysis. As GWAS meta-analysis of TB-BMD was also performed across five age stratum, MR was conducted in age-stratified manner.

Meanwhile, each SD decrease in genetically predicted blood LDL-C level was associated with 0.076 SD (95% CI: 0.042 to 0.111; P=1.20×10^−5^) increase in eBMD. Similar estimate was obtained in multivariable IVW model adjusting for beta estimates of HDL-C and triglycerides, as well as in sensitivity analyses of weighted median (0.065 SD increase in eBMD per SD decrease in LDL-C; 95% CI: 0.04 to 0.09; P=3.59×10^−7^) and MR-Egger method (0.052 SD increase in eBMD per SD decrease in LDL-C; 95% CI: −0.002 to 0.106; P=0.059). There was no evidence of pleiotropy for both phenotypes (MR-Egger intercept: 0.002; P=0.235 for TB-BMD; 0.002; P=0.257 for eBMD).

Reverse causation of BMD on LDL-C was also tested (Table 5). Each SD decrease in TB-BMD was associated with 0.035 SD increase in LDL-C (95% CI: 0.003 to 0.066; P=0.034) in univariable IVW analysis. Similar estimate with a wider confidence interval was obtained from weighted median method (0.043 SD increase in LDL-C per SD decrease in TB-BMD; 95% CI: −0.001 to 0.087; P=0.055) but insignificant association was suggested by MR-Egger method. There were no signs of pleiotropy (MR-Egger intercept: 0.002; P=0.486). The causal association remained significant in multivariable IVW analysis adjusting for HDL-C and triglycerides (0.037 SD increase in LDL-C per SD decrease in TB-BMD; 95% CI: 0.011 to 0.063; P=0.006). No reverse causation of eBMD on LDL-C was detected.

**Table 5.** Causal estimates for LDL-C (in SD) per 1 SD decrease of BMD

| **Mendelian Randomization approach to evaluate causal effects of TB-BMD on LDL-C** |  |  |  |  |
| --- | --- | --- | --- | --- |
| **Analysis** | **Beta** | **95% CI** |  | **P-value** |
|  |  | **Lower** | **Upper** |  |
| 68 SNPs |  |  |  |  |
| Conventional IVW | 0.035 | 0.003 | 0.066 | 0.034 |
| Multivariable IVW | 0.037 | 0.011 | 0.063 | 0.006 |
| Weighted median | 0.043 | -0.001 | 0.087 | 0.055 |
| MR-Egger | 0.004 | -0.087 | 0.095 | 0.927 |
| MR-Egger intercept | 0.002 | -0.003 | 0.006 | 0.486 |
| **Mendelian Randomization approach to evaluate causal effects of eBMD on LDL-C** |  |  |  |  |
| **Analysis** | **Beta** | **95% CI** |  | **P-value** |
|  |  | **Lower** | **Upper** |  |
| 221 SNPs |  |  |  |  |
| Conventional IVW | 0.025 | -0.007 | 0.058 | 0.128 |
| Multivariable IVW | 0.016 | -0.016 | 0.049 | 0.326 |
| Weighted median | 0.023 | -0.013 | 0.057 | 0.211 |
| MR-Egger | 0.006 | -0.065 | 0.077 | 0.867 |
| MR-Egger intercept | 0.001 | -0.002 | 0.004 | 0.547 |

For HDL-C, no significant association was observed with both BMD phenotypes. For triglycerides, univariable IVW analysis showed that 1 SD decrease was causally associated with 0.064 SD increase in TB-BMD (95% CI: 0.011 to 0.118; P=0.019). The causal association remained significant in multivariable IVW analyses adjusting for beta estimates of LDL-C and HDL-C. There was null causal association between triglycerides and eBMD.

### Drug target analysis using genetic proxies of LDL-C-lowering drugs

The effects of LDL-C-lowering drugs on BMD were examined by IVW analysis of their genetic proxies (Table 6). Statistical significance was detected for statins’ proxies where 1 SD decrease in genetically predicted decrease in LDL-C was associated with 0.18 SD increase in TB-BMD (95% CI: 0.044 to 0.316; P=9.600×10^−3^). Similar significant association was observed for eBMD (estimate: 0.143; 95% CI: 0.062 to 0.223; P=5.165×10^−4^). Genetic proxies for other LDL-C-lowering drugs did not show significant association with TB-BMD and eBMD.

**Table 6.** Causal estimates for BMD or fracture (in SD) per 1 SD decrease of LDL-C caused by LDL-C-lowering drugs

| Gene encoding<br>molecular target of<br>LDL-C-lowering<br>therapy | Beta | 95% CI |  | P-value |
| --- | --- | --- | --- | --- |
|  |  | Lower | Upper |  |
| Mendelian Randomization approach to evaluate the LDL-C-lowering effects on TB-BMD |  |  |  |  |
| HMGCR | 0.180 | 0.044 | 0.316 | 9.600 x 10<sup>-3</sup> |
| NPC1L1 | 0.080 | -0.138 | 0.298 | 0.472 |
| PCSK9 | 0.006 | -0.107 | 0.118 | 0.919 |
| ABCG5/G8 | 0.037 | -0.113 | 0.187 | 0.628 |
| LDLR | 0.018 | -0.064 | 0.101 | 0.667 |
| Mendelian Randomization approach to evaluate the LDL-C-lowering effects on eBMD |  |  |  |  |
| HMGCR | 0.143 | 0.062 | 0.223 | 5.165 x 10<sup>-4</sup> |
| NPC1L1 | -0.051 | -0.176 | 0.074 | 0.427 |
| PCSK9 | 0.039 | -0.011 | 0.089 | 0.125 |
| ABCG5/G8 | 0.037 | -0.049 | 0.123 | 0.396 |
| LDLR | 0.024 | -0.022 | 0.070 | 0.305 |
| Mendelian Randomization approach to evaluate the LDL-C-lowering effects on fracture |  |  |  |  |
| HMGCR | -0.090 | -0.285 | 0.104 | 0.363 |

### MR and drug target analysis – LDL-C and fracture

Causal association between LDL-C and fracture was evaluated. Null association was observed in univariable and multivariable MR analyses adjusted for HDL-C, triglycerides and eBMD. In view of statins’ LDL-C-lowering effects on increased BMD, drug target analysis was also performed on fracture. Null causal association was detected for genetic proxies of statin therapy on fracture (Table 6).

### MR analysis–TB-BMD, eBMD and CAD

The effects of TB-BMD and eBMD on CAD were assessed by univariable and multivariable MR adjusted for blood lipid levels, diabetes and BMI (Table 7). Null association was observed for TB-BMD on CAD. For eBMD, 1 SD decrease was associated with 5.1% reduced risk of CAD (odds ratio: 0.949; 95% CI: 0.903 to 0.998; P=0.042) in univariable IVW analysis. However, the association became insignificant in sensitivity analyses, and after taking risk factors (including blood LDL-C, HDL-C and triglycerides levels, diabetes and BMI) into account in multivariable analysis. Reverse causation of CAD on BMD was also tested but no significant association was observed.

**Table 7.** Causal estimates for CAD per 1 SD decrease of BMD

| <u>Mendelian Randomization approach to evaluate causality of TB-BMD on CAD</u> |  |  |  |  |
| --- | --- | --- | --- | --- |
| 68 SNPs |  |  |  |  |
| Analysis | Odds Ratio | 95% CI |  | P-value |
|  |  | Lower | Upper |  |
| Univariable MR analysis |  |  |  |  |
| Conventional IVW | 0.963 | 0.912 | 1.017 | 0.180 |
| Weighted median | 0.969 | 0.906 | 1.037 | 0.368 |
| MR-Egger | 1.059 | 0.911 | 1.231 | 0.453 |
| MR-Egger intercept | 0.995 | 0.987 | 1.003 | 0.186 |
| Multivariable IVW |  |  |  |  |
| LDL-C, HDL-C, triglycerides, diabetes and BMI as related risk factors | 0.980 | 0.925 | 1.039 | 0.504 |
| <u>Mendelian Randomization approach to evaluate causality of eBMD on CAD</u> |  |  |  |  |
| 214 SNPs |  |  |  |  |
| Analysis | Odds ratio | 95% CI |  | P-value |
|  |  | Lower | Upper |  |
| Univariable MR analysis |  |  |  |  |
| Conventional IVW | 0.949 | 0.903 | 0.998 | 0.042 |
| Weighted median | 0.973 | 0.921 | 1.027 | 0.320 |
| MR-Egger | 0.966 | 0.869 | 1.075 | 0.529 |
| MR-Egger intercept | 0.999 | 0.995 | 1.003 | 0.709 |
| Multivariable IVW |  |  |  |  |
| LDL-C, HDL-C, triglycerides, diabetes and BMI as related risk factors | 0.962 | 0.916 | 1.011 | 0.123 |

## Discussion

This study confirmed the association between LDL-C and BMD in two independent cohorts in epidemiological observation analyses and provides high-level evidence that a decrease in blood LDL-C level was causally associated with increased TB-BMD and eBMD. Whereas, decrease in TB-BMD, but not eBMD, was causally associated with increased LDL-C levels. The estimates derived from MR analyses were in line with the results from epidemiological observation analyses. MR analyses using genetic proxies of statin therapy consistently demonstrated that statins’ LDL-C-lowering effect was causally associated with increased TB-BMD and eBMD. We also examined the causal association between LDL-C and fracture, but null association was observed. There was insufficient evidence to prove the causation of BMD on CAD.

Our LD score regression analysis demonstrated moderate and positive genetic correlation between TB-BMD and eBMD. TB-BMD and eBMD are measured at different skeletal sites. eBMD is a quick and relatively inexpensive estimate of BMD, but it is not a proxy of DXA-measured BMD, which is the gold standard in the clinical management of osteoporosis. Notably, Kemp *et al*. reported that six eBMD-associated loci had opposite directions of effects when compared with GWAS of DXA-measured BMD^9^. DXA-measured BMD at sites prone to fracture (femoral neck and lumbar spine) has strong and positive genetic correlation with TB-BMD (r>0.9)^19^ but is just moderately correlated with eBMD (r=0.5–0.6) ^9^. TB-BMD is clinically more important than eBMD. We therefore investigated TB-BMD and eBMD as two different phenotypes in MR analyses. Importantly, LDL-C was shown to be associated with both TB-BMD and eBMD. Such robust evidence suggests that LDL-C is a causal factor involved in bone metabolism.

There has been controversy on the role of LDL-C in bone metabolism. Previous observational studies showed inconsistent association between LDL-C and BMD^3–7,10^, which may arise from unmeasured confounding and presence of reverse causation. As MR better accounts for these biases, our study, which cross-validated results of epidemiological observation analyses and MR, should provide a more reliable casual inference that reduction in LDL-C level is associated with increased BMD. It is worth-noting that power for the MR analysis of LDL-C on age-stratified TB-BMD is below 40% due to the small sample size in sub-groups (Table 1). The null association in sub-group analyses is likely attributed to inadequate power. Causal linkage between LDL-C level and life-course TB-BMD should be re-visited when future GWAS with larger age-stratified subgroups becomes available. In addition, our bi-directional MR analysis suggested that TB-BMD, but not eBMD, played a negative causal role in blood LDL-C level. Taken together, our study suggests a positive feedback loop between bone and lipid metabolism. This reinforces the role of bone in feedback control of energy homeostasis, and the concept of mutual regulation of bone and energy metabolism^35^.

We demonstrated that the LDL-C-lowering effect of statins was causally associated with increased TB-BMD and eBMD. In a recent meta-analysis investigating the effect of statins on BMD in both RCT and cohort studies, use of statins was significantly associated with increased BMD at lumbar spine (standardized mean difference [SMD]: 0.20; P=0.002), marginally significant at total hip (SMD: 0.18; P=0.05), but statistically insignificant at femoral neck (SMD: 0.08; P>0.05)^13^. Notably, only two small RCTs (total N=138) were included in the analysis, thus the significant result was mainly driven by the observational studies. In fact, null association with BMD was observed in the subgroup analysis of RCTs only. Another meta-analysis of RCTs, including the two small RCTs aforementioned and five other RCTs of larger sample size (N=27,754), suggested that statin use significantly increased BMD^14^. With consistent evidence from both observational and MR analyses, this study suggested causal inference of statin use on increased BMD.

Although causality could be inferred through MR, whether the finding could be translated clinically is unknown. Our findings suggested that lifelong genetic exposure to each SD decrease in LDL-C will increase TB-BMD by 0.038 SD or eBMD by 0.076 SD, but whether reducing LDL-C by 1 SD using pharmacological agent (e.g. statins) would lead to increase in eBMD / TB-BMD by the same amount is unknown, and such effect could not be tested in our observational cohorts due to the unavailability of pharmacological data. The effect of statins on LDL-C reduction was approximately 1.07mmol/L per year^36^ (~1.13 SD based on NHANES III data). In a meta-analysis of RCTs, statin treatment for one year was associated with 0.03g/cm^2^ increase in BMD (95% CI: 0.006, 0.053; P☐<☐0.001)^14^, which was approximately equivalent to 0.2 SD based on NHANES III data. Thus, it seems that reduction of LDL-C by statins may have an additional beneficial effect on BMD. These findings suggested the potential beneficial effects of statins on bone health, in addition to its protective role in cardiovascular diseases. As genetic predisposition to lower blood LDL-C levels is associated with increase in BMD, reduction of LDL-C level is therefore a common goal for the management of osteoporosis and CAD, the two prevalent diseases associated with increased immobility, morbidity and mortality.

Animal and cell studies provided mechanistic explanation regarding the role of LDL-C on bone metabolism. High cholesterol diet reduced BMD in mouse model^37^, probably via increased osteoclastogenesis^37,38^. On the other hand, statins target the mevalonate pathway, which is also the target of nitrogen-containing bisphosphonates (N-BPs), the first-line therapy for osteoporosis. Treatment of statins reduced bone loss via reduced osteoclastogenesis in ovariectomized rat^39^ and promoted osteoblast differentiation in ovariectomized rabbits^40^ respectively. As statins were reported to decrease bone resorption by inhibiting osteoclast differentiation and osteoblast apoptosis downstream of the mevalonate pathway^41^, this additional effect on bone resorption may explain why the reduction of LDL-C by statins may have an additional beneficial effect on BMD. While the mevalonate pathway is mainly responsible for cholesterol biosynthesis, the isoprenoid lipids produced are essential for prenylation and activation of small GTPases, which play a crucial role in the regulation of osteoclast morphology^42^. Studies reported that patients treated with intravenous N-BPs had a significant reduction in LDL-C level^43–46^ although studies examining patients treated with oral N-BP gave inconsistent results^47^. Use of N-BPs was also associated with lower risk of cardiovascular mortality and incident myocardial infarction in hip fracture patients^48^. Mevalonate pathway is the possible link between bone and lipid metabolism.

Our MR analyses demonstrated that decrease in LDL-C level was causally associated with increase in BMD but not fracture. Although LDL-C is a causal factor of BMD regulation, other non-bone related factors are also important in predicting fracture, such as muscle strength and risk of falls. These risk factors might explain the missing link between LDL-C level and fracture. Similarly, MR analyses suggested that the LDL-C-lowering effect of statins was causally associated with increased BMD, but it had null association with fracture. Meanwhile, statin use was found associated with reduced osteoporotic fracture in a meta-analysis of RCTs and observational studies^13^ though the finding was postulated to be confounded by healthy adherer effect^15^. The discrepancy may be explained by the fact that genetic proxies of statins just account for its LDL-lowering effects but could not fully cover its mechanism of action involving osteoblast differentiation and reduced osteoclastogenesis.

On the other hand, our MR analyses showed that decrease in triglycerides was causally associated with increase in TB-BMD in both univariable and multivariable IVW analyses. Yet, null association was observed for sensitivity analyses, which may be attributed to the low statistical power (25%).

Regarding the causal relationship between BMD and CAD, a MR study showed that increase in genetically-instrumented eBMD was significantly associated with higher risk of CAD^12^. It was inconsistent with the findings from a recent meta-analysis which showed that lower BMD was associated with a higher risk of CAD^11^. In our study, we showed that the effect of eBMD on CAD was indirect and it could be confounded by blood lipids levels^49^, diabetes^50,51^ and BMI^49^, which all shared common pathophysiological pathways with bone metabolism. In addition, our MR analyses suggested TB-BMD had a null effect on CAD. Insufficient evidence was present to prove the causal relation between BMD and CAD.

The present study has several strengths. Firstly, the epidemiological observation analyses were conducted in two independent cohorts: Mexican Americans, hispanic and non-hispanic in the U.S. population from the NHANES III, and southern Chinese from the HKOS. The consistent results across the two cohorts supported that LDL-C was negatively associated with BMD irrespective of ethnicities and living styles. Secondly, independent genetic instruments selected for the MR analyses were obtained from large-scale GWAS with stringent thresholds for quality-control and association analysis. The strength of combined genetic instruments in each MR analysis was assessed by the F-statistic, which ranged from 3,213.67 to 26,183.12. The high F-statistic indicated a lower chance of weak instrument bias. We had adequate statistical power to detect a causal effect (over 80%) for 11 out of the 13 main MR analyses conducted (Table 1), assuming the causal beta coefficient is the same as the beta estimate obtained from epidemiological observational studies for continuous outcome. Thirdly, in evaluating the causality of blood LDL-C level on TB-BMD and eBMD, two-sample MR analysis was adopted and there was no overlap between the samples of blood lipids and BMD. Therefore, over-fitting of the findings was avoided. The estimated effects derived by MR approach was similar to that derived from the epidemiological observation analysis conducted in two cohorts. Consistent results were also observed in various sensitivity analyses, suggesting that the results were robust and confounding factors were unlikely to explain the observed associations.

This study also has limitations. In the epidemiological observation study, BMD at femoral neck and lumbar spine were investigated as these sites were known to be prone to fracture. Whereas, TB-BMD was investigated in MR analysis as the GWAS meta-analysis^19^ contains the largest sample for DXA-derived BMD to date and it is closely correlated with BMD at lumbar spine and femoral neck (r>0.9)^19^. Meanwhile, eBMD was examined as an alternative BMD phenotype with a large GWAS dataset of over 140,000 participants. The large sample size of the TB-BMD and eBMD datasets enable our study to have sufficient statistical power to detect genuine causal effects. MR does not require prior understanding on the functions of the genetic instruments and how they influence the risk factors. It is possible that the genetic instruments may have an indirect effect on the outcome via a pathway that does not involve the risk factor of interest (horizontal pleiotropy). Our sensitivity analyses showed no evidence for unbalanced horizontal pleiotropy, though it cannot be ruled out unequivocally. Regarding the MR analysis of eBMD on CAD, both datasets included participants from UK Biobank and there were likely overlapping samples. With causal association detected, bias would be present in the direction of the confounded association and the net bias would rely on the degree of overlap which could not be accounted for^52^.

## Conclusion

In conclusion, the current study provided strong evidence that genetic predisposition to lower blood LDL-C levels was associated with increase in BMD. TB-BMD also had a negative causal role on LDL-C level. MR analysis using the genetic proxies of statin therapy demonstrated that statins’ LDL-C-lowering effects could improve BMD. Insufficient evidence was available to support the causation of LDL-C on fracture, and BMD on CAD.

## References

1. Farhat GN, Cauley JA. The link between osteoporosis and cardiovascular disease. Clin Cases Miner Bone Metab. 2008;5(1):19–34.

2. Ference BA, Ginsberg HN, Graham I, et al. Low-density lipoproteins cause atherosclerotic cardiovascular disease. 1. Evidence from genetic, epidemiologic, and clinical studies. A consensus statement from the European Atherosclerosis Society Consensus Panel. Eur Heart J. 2017;38(32):2459–2472.

3. Kuipers AL, Miljkovic I, Evans R, Bunker CH, Patrick AL, Zmuda JM. Optimal serum cholesterol concentrations are associated with accelerated bone loss in African ancestry men. Osteoporosis international: a journal established as result of cooperation between the European Foundation for Osteoporosis and the National Osteoporosis Foundation of the USA. 2016;27(4):1577–1584.

4. Martin-Gonzalez C, Gonzalez-Reimers E, Quintero-Platt G, et al. Lipid profile and bone mineral density in heavy alcoholics. Clinical nutrition. 2017.

5. Li S, Guo H, Liu Y, et al. Relationships of serum lipid profiles and bone mineral density in postmenopausal Chinese women. Clinical endocrinology. 2015;82(1):53–58.

6. Jeong IK, Cho SW, Kim SW, et al. Lipid profiles and bone mineral density in pre- and postmenopausal women in Korea. Calcified tissue international. 2010;87(6):507–512.

7. Garg MK, Marwaha RK, Tandon N, Bhadra K, Mahalle N. Relationship of lipid parameters with bone mineral density in Indian population. Indian journal of endocrinology and metabolism. 2014;18(3):325–332.

8. Andersen L, Ibarra J, Andersen R. Current familial hypercholesterolemia diagnostic criteria underdiagnose APOB mutations: Lessons from the Amish community. J Clin Lipidol. 2016;10(2):443–444.

9. Kemp JP, Morris JA, Medina-Gomez C, et al. Identification of 153 new loci associated with heel bone mineral density and functional involvement of GPC6 in osteoporosis. Nat Genet. 2017;49(10):1468–1475.

10. Chen YY, Wang WW, Yang L, Chen WW, Zhang HX. Association between lipid profiles and osteoporosis in postmenopausal women: a meta-analysis. European review for medical and pharmacological sciences. 2018;22(1):1–9.

11. Veronese N, Stubbs B, Crepaldi G, et al. Relationship Between Low Bone Mineral Density and Fractures With Incident Cardiovascular Disease: A Systematic Review and Meta-Analysis. Journal of bone and mineral research: the official journal of the American Society for Bone and Mineral Research. 2017;32(5):1126–1135.

12. Gan W, Clarke RJ, Mahajan A, et al. Bone mineral density and risk of type 2 diabetes and coronary heart disease: A Mendelian randomization study. Wellcome Open Res. 2017;2:68.

13. An T, Hao J, Sun S, et al. Efficacy of statins for osteoporosis: a systematic review and meta-analysis. Osteoporosis international: a journal established as result of cooperation between the European Foundation for Osteoporosis and the National Osteoporosis Foundation of the USA. 2017;28(1):47–57.

14. Wang Z, Li Y, Zhou F, Piao Z, Hao J. Effects of Statins on Bone Mineral Density and Fracture Risk: A PRISMA-compliant Systematic Review and Meta-Analysis. Medicine. 2016;95(22):e3042.

15. Donzelli A, Battaggia A, Schivalocchi A. Statin use does not protect from fractures: the healthy adherer effect is a plausible explanation in observational studies. Osteoporosis international: a journal established as result of cooperation between the European Foundation for Osteoporosis and the National Osteoporosis Foundation of the USA. 2017;28(9):2739–2740.

16. Sertbas Y, Ersoy U, Ayter M, Gultekin Tirtil F, Kucukkaya B. Ezetimibe effect on bone mineral density and markers of bone formation and resorption. J Investig Med. 2010;58(2):295–297.

17. Wadhera RK, Steen DL, Khan I, Giugliano RP, Foody JM. A review of low-density lipoprotein cholesterol, treatment strategies, and its impact on cardiovascular disease morbidity and mortality. J Clin Lipidol. 2016;10(3):472–489.

18. Davies NM, Holmes MV, Davey Smith G. Reading Mendelian randomisation studies: a guide, glossary, and checklist for clinicians. BMJ. 2018;362:k601.

19. Medina-Gomez C, Kemp JP, Trajanoska K, et al. Life-Course Genome-wide Association Study Meta-analysis of Total Body BMD and Assessment of Age-Specific Effects. Am J Hum Genet. 2018;102(1):88–102.

20. Willer CJ, Schmidt EM, Sengupta S, et al. Discovery and refinement of loci associated with lipid levels. Nat Genet. 2013;45(11):1274–1283.

21. Trajanoska K, Morris JA, Oei L, et al. Assessment of the genetic and clinical determinants of fracture risk: genome wide association and mendelian randomisation study. Bmj. 2018;362:k3225.

22. Nelson CP, Goel A, Butterworth AS, et al. Association analyses based on false discovery rate implicate new loci for coronary artery disease. Nat Genet. 2017;49(9):1385–1391.

23. Scott RA, Scott LJ, Magi R, et al. An Expanded Genome-Wide Association Study of Type 2 Diabetes in Europeans. Diabetes. 2017;66(11):2888–2902.

24. Yengo L, Sidorenko J, Kemper KE, et al. Meta-analysis of genome-wide association studies for height and body mass index in approximately 700000 individuals of European ancestry. Hum Mol Genet. 2018.

25. Bulik-Sullivan B, Finucane HK, Anttila V, et al. An atlas of genetic correlations across human diseases and traits. Nat Genet. 2015;47(11):1236–1241.

26. Bulik-Sullivan BK, Loh PR, Finucane HK, et al. LD Score regression distinguishes confounding from polygenicity in genome-wide association studies. Nat Genet. 2015;47(3):291–295.

27. Burgess S, Butterworth A, Thompson SG. Mendelian randomization analysis with multiple genetic variants using summarized data. Genetic epidemiology. 2013;37(7):658–665.

28. Burgess S, Thompson SG. Multivariable Mendelian randomization: the use of pleiotropic genetic variants to estimate causal effects. Am J Epidemiol. 2015;181(4):251–260.

29. Bowden J, Davey Smith G, Haycock PC, Burgess S. Consistent Estimation in Mendelian Randomization with Some Invalid Instruments Using a Weighted Median Estimator. Genetic epidemiology. 2016;40(4):304–314.

30. Bowden J, Davey Smith G, Burgess S. Mendelian randomization with invalid instruments: effect estimation and bias detection through Egger regression. Int J Epidemiol. 2015;44(2):512–525.

31. Yavorska OO, Burgess S. MendelianRandomization: an R package for performing Mendelian randomization analyses using summarized data. Int J Epidemiol. 2017;46(6):1734–1739.

32. Brion MJ, Shakhbazov K, Visscher PM. Calculating statistical power in Mendelian randomization studies. Int J Epidemiol. 2013;42(5):1497–1501.

33. Lotta LA, Sharp SJ, Burgess S, et al. Association Between Low-Density Lipoprotein Cholesterol-Lowering Genetic Variants and Risk of Type 2 Diabetes: A Meta-analysis. JAMA. 2016;316(13):1383–1391.

34. Do R, Willer CJ, Schmidt EM, et al. Common variants associated with plasma triglycerides and risk for coronary artery disease. Nat Genet. 2013;45(11):1345–1352.

35. Lee NK, Sowa H, Hinoi E, et al. Endocrine regulation of energy metabolism by the skeleton. Cell. 2007;130(3):456–469.

36. Cholesterol Treatment Trialists C, Baigent C, Blackwell L, et al. Efficacy and safety of more intensive lowering of LDL cholesterol: a meta-analysis of data from 170,000 participants in 26 randomised trials. Lancet. 2010;376(9753):1670–1681.

37. Pelton K, Krieder J, Joiner D, Freeman MR, Goldstein SA, Solomon KR. Hypercholesterolemia promotes an osteoporotic phenotype. The American journal of pathology. 2012;181(3):928–936.

38. Sanbe T, Tomofuji T, Ekuni D, Azuma T, Tamaki N, Yamamoto T. Oral administration of vitamin C prevents alveolar bone resorption induced by high dietary cholesterol in rats. Journal of periodontology. 2007;78(11):2165–2170.

39. Oxlund H, Andreassen TT. Simvastatin treatment partially prevents ovariectomy-induced bone loss while increasing cortical bone formation. Bone. 2004;34(4):609–618.

40. Zhou H, Xie Y, Baloch Z, Shi Q, Huo Q, Ma T. The effect of atorvastatin, 3-hydroxy-3-methylglutaryl coenzyme A reductase inhibitor (HMG-CoA), on the prevention of osteoporosis in ovariectomized rabbits. J Bone Miner Metab. 2017;35(3):245–254.

41. Ruan F, Zheng Q, Wang J. Mechanisms of bone anabolism regulated by statins. Biosci Rep. 2012;32(6):511–519.

42. Coxon FP, Rogers MJ. The role of prenylated small GTP-binding proteins in the regulation of osteoclast function. Calcified tissue international. 2003;72(1):80–84.

43. Adami S, Braga V, Guidi G, Gatti D, Gerardi D, Fracassi E. Chronic intravenous aminobisphosphonate therapy increases high-density lipoprotein cholesterol and decreases low-density lipoprotein cholesterol. Journal of bone and mineral research: the official journal of the American Society for Bone and Mineral Research. 2000;15(3):599–604.

44. Montagnani A, Gonnelli S, Cepollaro C, et al. Changes in serum HDL and LDL cholesterol in patients with Paget’s bone disease treated with pamidronate. Bone. 2003;32(1):15–19.

45. Gozzetti A, Gennari L, Merlotti D, et al. The effects of zoledronic acid on serum lipids in multiple myeloma patients. Calcified tissue international. 2008;82(4):258–262.

46. Gonnelli S, Caffarelli C, Tanzilli L, et al. Effects of intravenous zoledronate and ibandronate on carotid intima-media thickness, lipids and FGF-23 in postmenopausal osteoporotic women. Bone. 2014;61:27–32.

47. Caffarelli C, Montagnani A, Nuti R, Gonnelli S. Bisphosphonates, atherosclerosis and vascular calcification: update and systematic review of clinical studies. Clin Interv Aging. 2017;12:1819–1828.

48. Sing CW, Wong AY, Kiel DP, et al. Association of Alendronate and Risk of Cardiovascular Events in Patients With Hip Fracture. Journal of bone and mineral research: the official journal of the American Society for Bone and Mineral Research. 2018.

49. Palermo A, Tuccinardi D, Defeudis G, et al. BMI and BMD: The Potential Interplay between Obesity and Bone Fragility. Int J Environ Res Public Health. 2016;13(6).

50. Compston J. Type 2 diabetes mellitus and bone. J Intern Med. 2018;283(2):140–153.

51. Valderrabano RJ, Linares MI. Diabetes mellitus and bone health: epidemiology, etiology and implications for fracture risk stratification. Clin Diabetes Endocrinol. 2018;4:9.

52. Burgess S, Davies NM, Thompson SG. Bias due to participant overlap in two-sample Mendelian randomization. Genetic epidemiology. 2016;40(7):597–608.

